# Development of locomotion in low water exposure using sturgeon

**DOI:** 10.1101/2020.03.01.972257

**Authors:** Anshin. Asano-Hoshino, Hideyuki. Tanaka, Takashi. Nakakura, Toshiaki. Tsuji, Takuo. Mizukami

## Abstract

The evolution of early land vertebrates from aquatic forms of life was a biological milestone. The transition to land was accompanied with expectedly challenging physiological and morphological evolutionary hurdles. So far, fossil records have provided substantial information on the origin of quadrupedal locomotion. However, fossil evidence alone is insufficient to understand how the soft-tissue-dependent motor functions and locomotion were acquired and developed. In the present study, we focus on locomotion of the sturgeon, an extant primitive fish, as a new experimental model, to investigate behavioural plasticity. Their locomotion in low-water-level conditions was similar to an escape response in water, the C-start escape response, which is used by most fish and amphibian juveniles to avoid predation. Sturgeons were also found to have mastered rolling-over in response to low water levels, resulting in the improvement of their trunk-twisting action. Sturgeons acquired an efficient shift in their centroid, thereby improving their mobility. We hypothesise that the escape response triggered by environmental hazards drove the development of locomotion, which was accompanied by a variety of behaviours.

## Introduction

Animal locomotion is any of a variety of ways that animals use to move from one place to another. Forms of locomotion include, walking, running, jumping, swimming, and flying. About 20,000 species of fish live in the hydrosphere, including the sea, rivers, and lakes. In general, fish live a lifetime in the water, but some fish live in intertidal zones and tidal flats. These fish are constantly exposed to environmental changes compared to underwater. Therefore, these fishes have modified their behaviour and physiological functions suitable for the habitat (Nelson, 2006) In the Devonian period, 400 million years ago, the pectoral and pelvic fins of fish evolved into appendicular structures, enabling terrestrial locomotion. However, it remains unclear what kind of environment provoked the evolution of these appendicular structures and what kind of mechanisms contributed to the development of terrestrial locomotion (Goetz et al., 2015; Grande and Bemis, 1991; Gregory and Raven, 1941; Niedźwiedzki et al., 2010). Recent studies have reported some interesting results in morphological and behavioural approaches to understanding the origins of quadrupedal locomotion. King et al. (King et al., 2011) observed the underwater behaviour of lungfish, one of the few species of extant finned sarcopterygians, which showed locomotion similar to terrestrial walking. Other studies demonstrated that Polypterus, the extant fish closest to the common ancestor of actinopterygians and sarcopterygians, exhibited developmental and phenotypic plasticity when exposed to low water levels (Standen et al., 2014; Standen et al., 2016) Lungfish and Polypterus are both capable of spontaneously moving onto land in search for water and food and, thus, already seem to have terrestrial locomotive function (Du and Standen, 2017; Du et al., 2016; Graham et al., 2014). It is important, therefore, to focus on behavioural plasticity by using fsih that is always swimming in the water as new experimental models and analysis system to make inferences on the diverse processes that led to terrestrialisation. From an evolutionary perspective, we are interested in how unspecialised fishes that constantly swim in water locomote when exposed to different terrestrial environments. In this study, we experimentally exposed sturgeon to terrestrial environments, with the aim of visualisation and quantification of changes in its locomotion during the adaptation from aquatic to terrestrial environments; we also explored the mechanisms underlying this process..

## Materials and methods

### Experimental animals

We used Bester, a hybrid sturgeon between *Huso huso* and *Acipenser ruthenus*. Fujikin Corporation provided the fish. Reliable non-invasive sex determination of Bester is not anatomically possible; as a result, animals were kept and studied in unmarked mixed-sex groups. Animal growth was monitored by assessing the fish length and weight upon entry in the environments. All fish had been raised in fully aquatic environments (n = 42). The fish were divided into the aquatic environment (control, n = 10) and the terrestrial environment (treatment, n = 52) groups. We performed animal experiments in compliance with the regulation for animal experiments of Teikyo University.

### Rearing environments

Two 400-L and one 200-L overflow tanks were connected to a 200-L filtration tank, and used as a circulation/filtration system with a total water capacity of 1.2 m^3^. Each tank was connected to an airstone using a blower air pump for the septic tank (Nippon Denko, NIP-40L) with aeration. The sturgeons were fed 2–3 times a day with pellet-type feed (feed dedicated to carnivorous benthic fish, Kyorin, Hikari Crest Cat) suitable for their size that allows rapid consumption.

### Exposure to low water levels

The room temperature was set at 20 °C. An automated mist system (Foresta, Zero Plants) and a shower pipe (EHEIM) were used regularly, and the water quality at low water levels was ensured by constantly pouring water through 2-mm pores at the top of tanks. Since the sturgeon cannot eat during low-water-level conditions due to its suction-type mouth, 5 days a week were set at low-water-level, whereas the remaining 2 days at normal water level to allow feeding and swimming. Their body weights decreased during the low-water-level conditions, but recovered in the following 2 days, totally resulting in weight gains in a week.

### Measurements

Quantitative analyses on zero moment point (ZMP) during the exposure of sturgeons to low water levels were performed using the Haptic Cutting Board system and Digital Array system (Pressure Profile System Co.), which we developed in collaboration with Saitama University. This sensor enabled us to visualise the loads involved in the locomotion in more detail. We visualised the temporal changes in the positional information of a point, called the ZMP, and the dynamic components during the low-water-level locomotion in three-dimensional vectors (Totsu et al., 2015; Tsuji et al., 2009). In other words, ZMP is a dynamic barycentric centre, implying that biped locomotion could be achieved by applying evolutionary constraints and conditions that repeatedly forced ZMP to achieve locomotion on land.

### Image files

All behavioural sequences were filmed at frames 1080p/60fps using i-phone 6S (Apple) and full-HD HDC-TM750 Camera (Panasonic). The following picture was created by captured movie files.

## Results

Our observations of sturgeons swimming patterns under normal water conditions revealed that their pectoral fins do not contribute substantially to locomotion, whereas the trunks operate in a wavelike manner to drive forward movement (Figure 1A) (LINDSEY and CC, 1978; Wilga and Lauder, 1999). This observation indicates that the sturgeon has higher propulsion during underwater locomotion than the lungfish and the polypterus, which are benthic and generally remain stationary.

**Figure 1.**
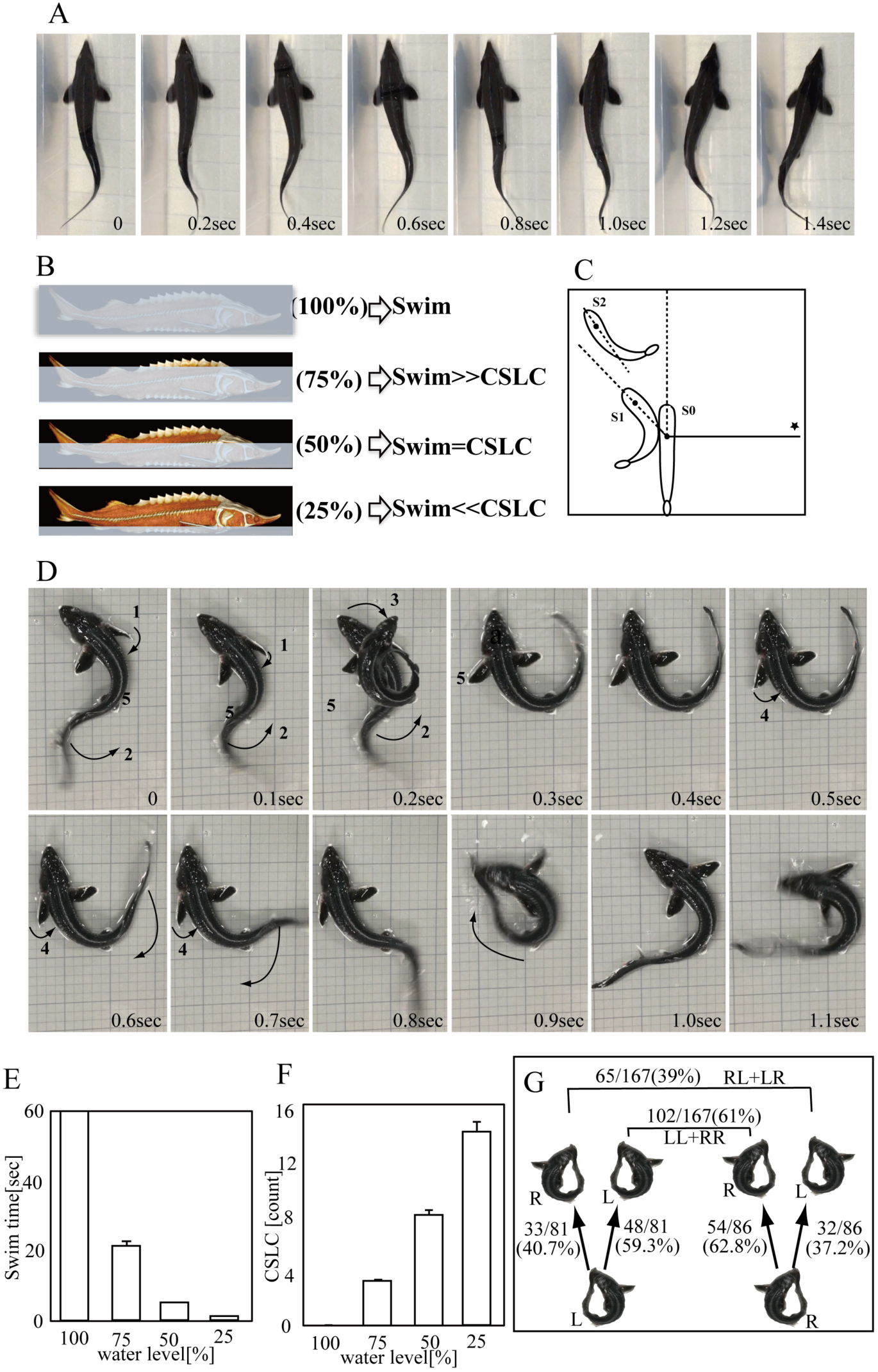
Changes in water-level-dependent locomotion of a sturgeon. (A) Swimming underwater. (B) Changes in locomotion at different water levels. (C) Schematic diagram of C-start escape response during underwater swimming. Numbers indicate the order of motions. S0: The point where the fish received a stimulus. S1: C-shaped escape behaviour. S2: The position of escape using a rebound. * Position of the sound source. (D) Sequential photographs of C-start-like crawling (CSLC) motion. (E) Swimming time in one minute at each water level. (F) CSLC measurement in one minute at each water level. (G) Schematic diagram of the rate of C-start escape response on the land.

The sturgeons could thrive even when the water level was lowered to the eye level. Under 100% water level (whole-body immersion), sturgeons only exhibited swimming with an S-shaped wavelike movement of the trunk. When the water was lowered to 75%, crawling movements with pectoral fins, folded on alternate sides, were often observed. At 50% water level, the frequency of crawling increased, and when it was further lowered to 25%, we observed crawling at a higher frequency. In particular, Polypterus showed both behavioural and quantitative changes in terms of locomotion (Figures 1B, 1D, 1E, and 1F). Thus, our results indicate that the locomotion of the sturgeon underwater is swimming with an S-shaped wavelike movement of the trunk, and it switches to crawling depending on the water level (Figures 1E and 1F). This switching may be mediated by the perceived decrease in propulsion, as well as the changes in sound, gravity, and water flow.

We carefully observed the crawling movement and found that the sturgeons showed movements similar to a reflex response, called the C-start escape response. Fish and amphibians share this response when they perceive hazardous stimuli (Figure 1D and Supplementary Material, Movie1_)_. We named this response the “C-start-like crawling” (from now on referred to as CSLC). The C-start escape response is a sudden escape response behaviour that the juveniles of most fish and amphibians (e.g., tadpoles) exhibit in response to “fear” stimuli, such as sounds and waves underwater. The response allows them to accelerate and escape quickly and involves a C-shape contraction of the side muscles on the opposite side from the stimuli, and accompanied by swings of the caudal fin (Figure 1C) (Domenici and Blake, 1997; Wassersug, 1989). A peculiar single movement is observed when fish show escape response behaviours underwater(Watanabe et al., 2014), whereas the sturgeons exposed to low water levels in our study used sequential CSLC for locomotion. The reflex responses appeared repeatedly on the same side (61%), unlike terrestrial walking, which involves alternating movements (39%) (Figure 1G). Sturgeons with body lengths of 20 cm had an average moving distance of 3.49 ± 0.78 cm and 106.24 ± 9.11° at a time (Supplementary Material figure S2). We observed CSLC in all fish (n = 40) immediately after exposure to stimuli. These results suggest that many kinds of fish are capable of terrestrial locomotion once adapted to the exposure to the atmosphere. Similar experiments on young sturgeons (1–2 weeks old) and Japanese catfish revealed C-start escape response behaviours immediately after exposure to stimuli (Supplementary Material figure S1).

We also checked sequential ventral photographs of CSLC motion. In this experiment, CSLC was viewed from the ventral side of the fish on a glass plate; interestingly, we observed that the pelvic fins were in contact with the glass plate when the sturgeon moved forward using CSLC. On the other hand, when the sturgeon bent its body by moving its tail and head, its pelvic fins were not in contact with the glass (Figure 2).

**Figure 2.**
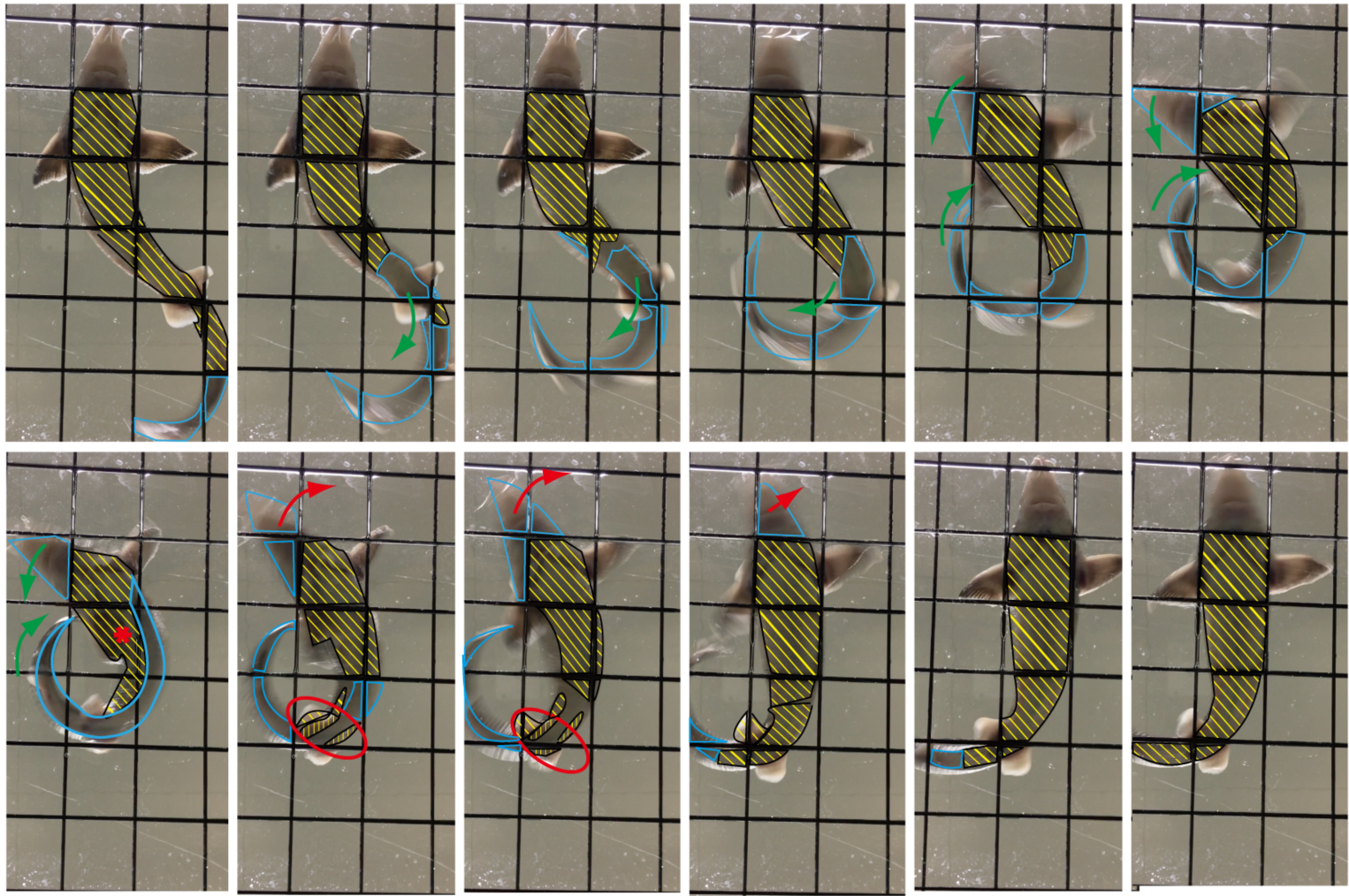
Sequential ventral photographs of C-start-like crawling (CSLC) motion. Observation of the bottom surface of sturgeons during CSLC. The area surrounded by a light blue line indicates the area that is not in contact with the ground. The yellow shaded region is in contact with the ground. The red circle indicates the loaded pelvic fin region. * Position of ZMP. The green arrows and red arrows indicate the direction of lateral bending.

Based on the measurements of the surface pressure distribution during locomotion under low-water-level conditions, we found that the sturgeons could only apply weak loads to the entire lower surface of their trunks with CSLC (Figure 3A). On the other hand, the sturgeons that acquired the twisting action before producing long-distance movements could apply strong and timely local loads, with the heaviest load applied via the lateral bending C-shape. Interestingly, we found that the sturgeons mechanically applied the loads not only to the pectoral fin region and the thoracic appendage but also to the pelvic fin region and the abdominal appendage (Figure 3B). These results indicate that rearing in low-water-level conditions enabled CSLC-mediated load control during locomotion using not only pectoral but also pelvic fins. These results, together with a previous report showing that the escape route changed when pelvic fins were removed, also suggest that the pelvic fins act as the “brake,” and is the turning axis for the C-start escape response underwater (Kawabata et al., 2016; Standen, 2008) and that the pectoral and pelvic fins are involved in the initial terrestrial locomotion.

**Figure 3.**
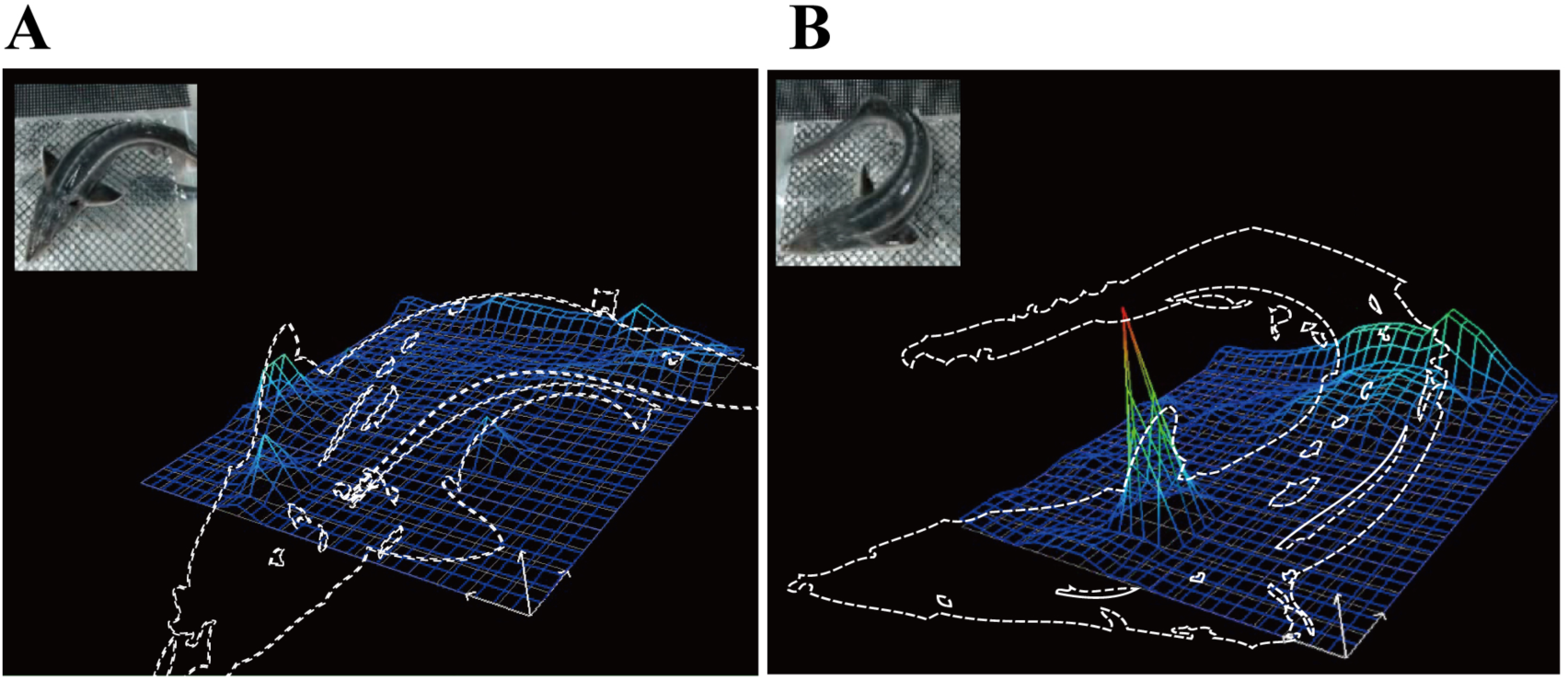
Detection of floor-reaction forces in the CSLC of sturgeons. (A) A weak load was detected under some regions of whole sturgeons. (B) A temporary strong load was detected under the pectoral and pelvic fin region of sturgeons.

After exposure to low water levels, the sturgeons were unable to roll over to a normal posture for several hours. However, after rearing them in this environment for a few days, we found that most of them had mastered the ability to roll over to a normal posture while repeating lateral bending, which is normally used to swim underwater (Figure 4A). The rolling-over indicated the development of a trunk-twisting action (Figure 4B). Consequently, when we tried to roll them over, by applying resistance, they could recover their normal posture, often assuming a posture with their pectoral fins fully expanded when lying in the prone position. In addition, fish were able to move by jumping, resulting in dramatically longer movements (Figures S4 and 6G). Thus, the sturgeons distinctively adapt to the new environment and develop their motor functions, through enhanced trunk-twisting action. These adaptations generally occurred within the first 10 days after the exposure to low water levels (Figure 4C).

**Figure 4.**
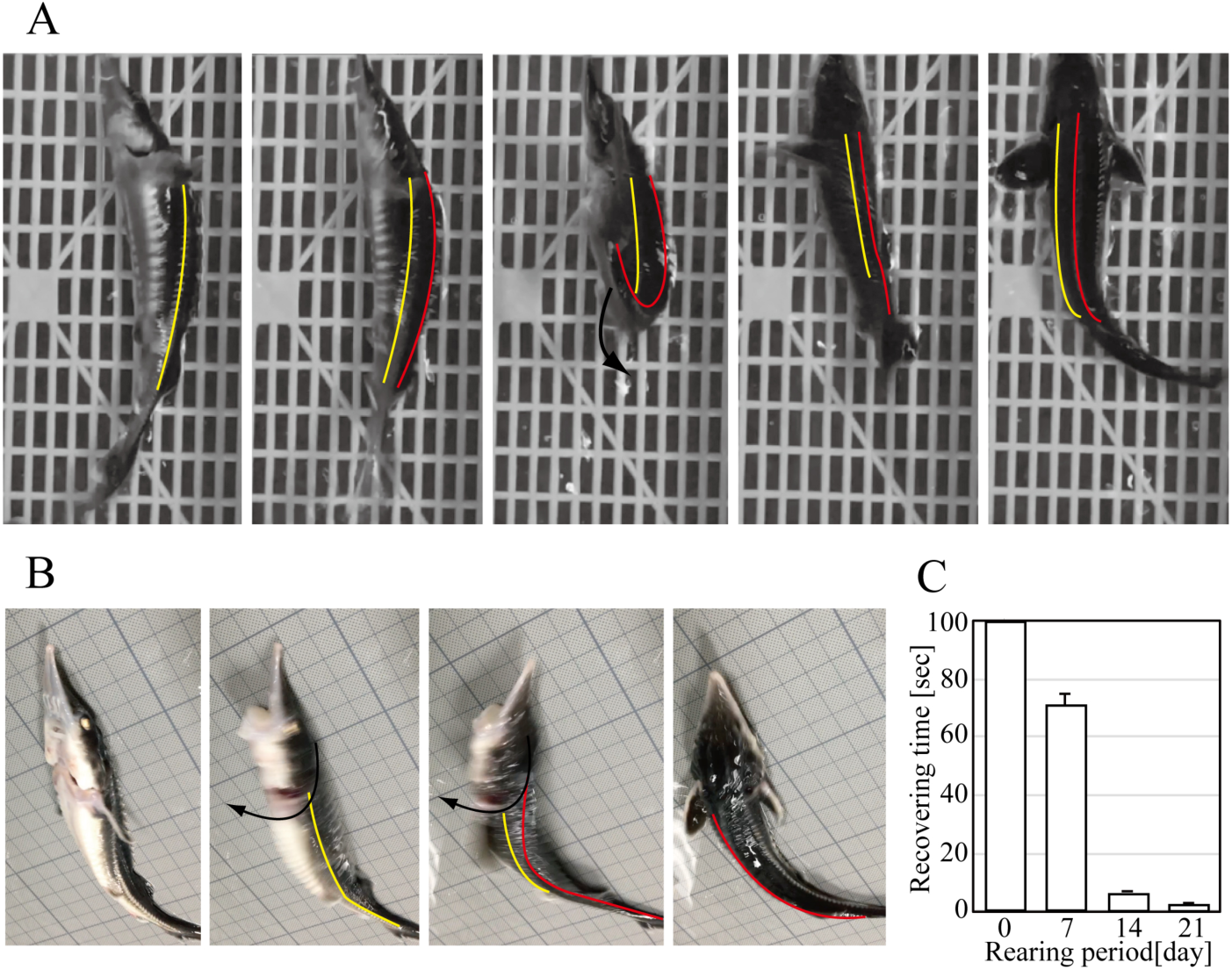
Development of locomotion. (A) Rolling-over using C-start escape response. (B) Rolling-over using twisting force. The red lines indicate the dorsal side of the trunk, and the yellow lines indicate the lateral side of the trunk. (C) The recovery time decreased depending on the exposure period.

**Figure 5.**
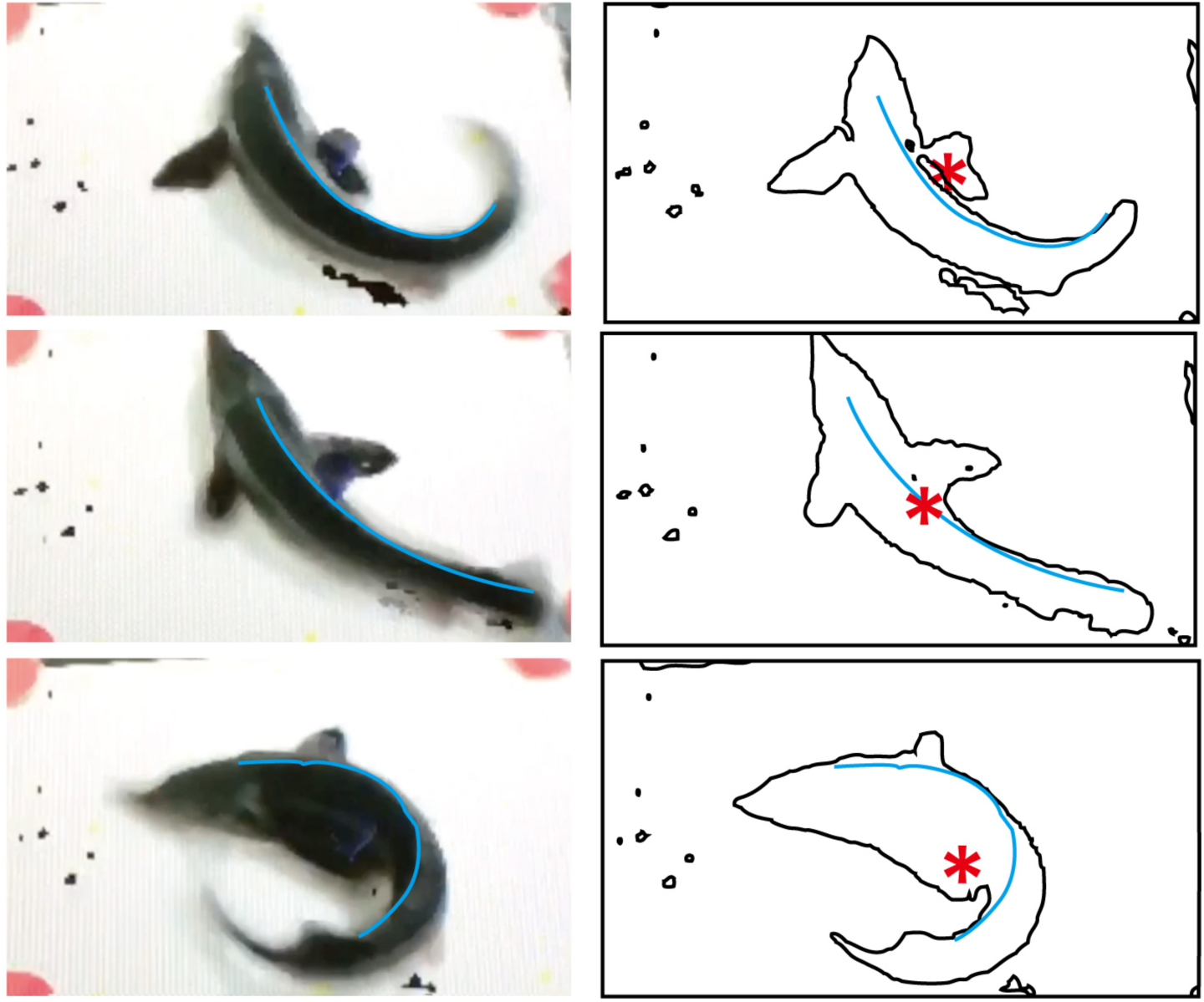
Changes in the barycentric centre and load in the locomotion process during exposure to low water levels. Detection of the position of the barycentric centre during CSLC locomotion. The left panel is the still image from the movie file. The right panel is the outline of the image in the left panel. * Position of ZMP.

**Figure 6.**
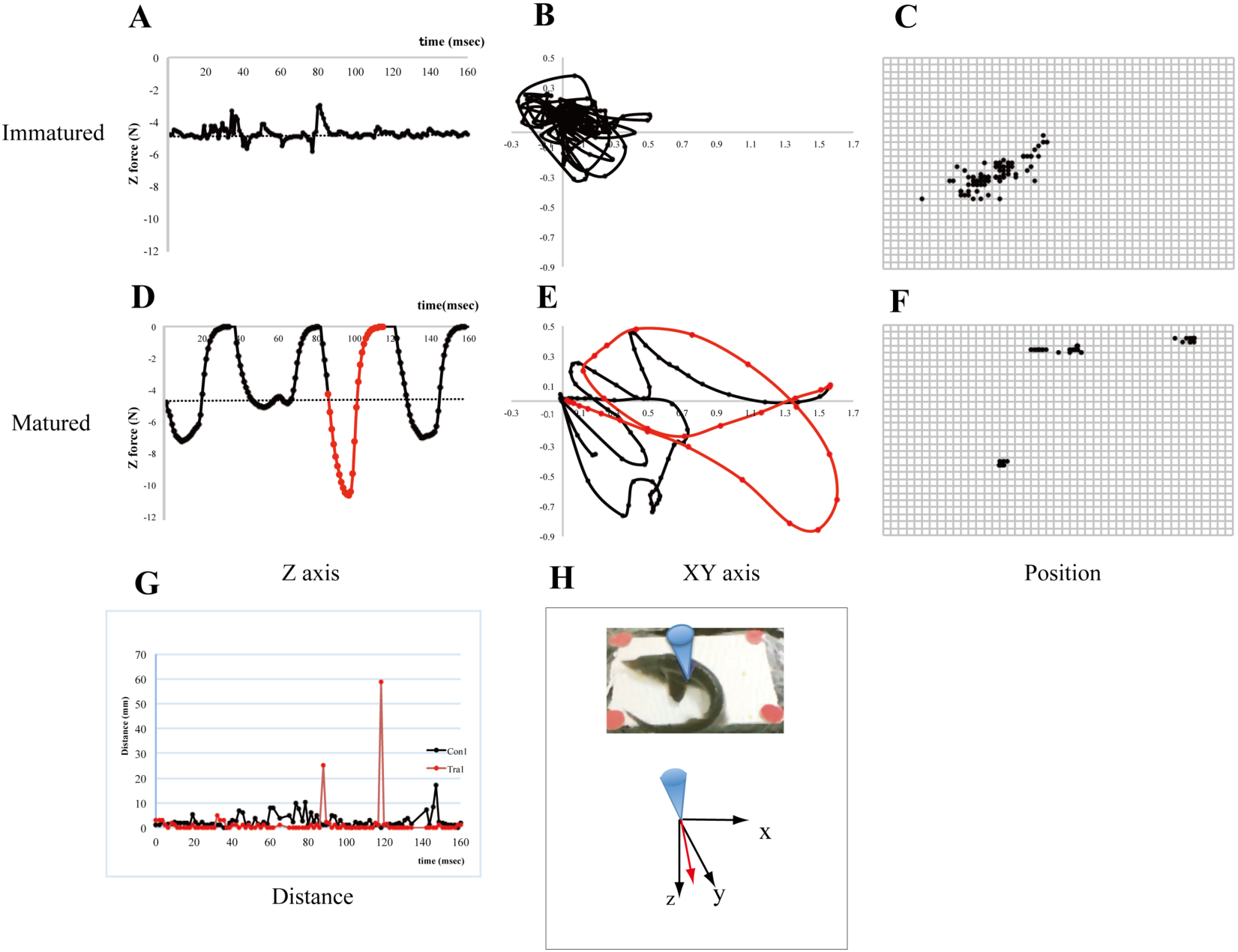
Measurement of the direction and magnitude of the force of ZMP in CSLC using a tactile sensor. After long-term exposure to low water levels, the sturgeons could apply more load in the vertical direction (A and D) and change the loads more in the horizontal direction (B and E), compared with the sturgeons immediately after exposure. The position of ZMP changed gradually with short moving distances in the sturgeons immediately after exposure (C and G), whereas the sturgeons after long-term exposure did not usually move much and changed their position significantly only when they moved (F and G). Changes in the vertical loads in CSLC at the ZMP point over time (A and D). The dashed line indicates the level of its weight. (B) and (E): Temporal changes in the direction of horizontal forces in CSLC. (C) and (F): Distribution of ZMP in CSLC. (G) Moved distance of ZMP. (H) Schematic diagram of the Haptic Cutting Board system device. The upper panel shows an immature locomotion group (A, B, and C) and the lower panel mature locomotion group (D, E, and F), Con: control group, Tra: low-water-level exposed group. The red lines in B and E indicate the same time point.

We observed that the locomotion pattern changed, and the moving distance extended during the development of sturgeon locomotion, which was enabled by an originally developed tactile sensor system (BARCLAY, 1946; Totsu et al., 2015; Tsuji et al., 2009). This system can help visualize and measure the direction and magnitude of forces applied to support surfaces on a millisecond scale. Our results using this system revealed that the position of the ZMP during the locomotion process by CSLC moved right and left slightly behind the pectoral fins in conjunction with trunk bending (Figure 5). This finding indicates that the pressure centre of the floor-reaction force is located close to the pectoral fin. Therefore, propulsion on the land mainly used the front load of the pectoral fin region.

Detailed quantitative analysis of CSLC in sturgeons immediately after exposure and after the acquisition of the ability to roll over, revealed that the former moved little by little with a distance of 10 mm or less (Figures 6C and 6G), while the latter stayed stationary without locomotion but could move long distances instantaneously during locomotion by CSLC (Figures 6F and 6G). In comparison with the sturgeons immediately after exposure to low water levels, the vector of ZMP in the sturgeons after acquiring the ability to roll over generated loads 2.3 times (Figures 6B and 6E) and 1.9 times (Figures 6A and 6D) more efficiently in the horizontal (X and Y axes) and vertical (Z axis) directions. These results culminate with the observed increase (× 3.3) in the moving distance (Figure 6G). These results indicate that the locomotion of sturgeons under low-water-level conditions was improved by effectively controlling the load onto the ground, as a result of a stronger twisting force around the body axes, mediated by the ability to roll over.

## Discussion

In this study, we visualised fish locomotion by experimentally exposing sturgeons to low water levels. Since CSLC is the escape response typical to all fish in water, many kinds of fish could have potentially used it for locomotion. The sturgeons appeared to become stationary a few minutes after exposure to low water levels, and subsequently, they moved and changed directions by continuously performing complicated and irregular CSLCs. The continuity of locomotion observed may reflect continuous reflex responses in response to constant stimuli due to environmental changes, such as gravity and sound, during the shift from aquatic to terrestrial environments. New-born zebrafish—which exhibit motor functions comparable to those of adult fish on the second day after fertilisation—reportedly show the continuous firing of C-start escape responses during the early stages, but later acquire single firing of Meissner’s corpuscle in association with the development of auditory inputs. In this experiment, the lateral line, a sensory organ that senses water flow in fish, as well as auditory sensations, could have influenced the sturgeons in various ways, leading to the development of terrestrial locomotion. Our mechanical analysis demonstrated that the change in locomotion over time allowed the sturgeon to acquire a rolling-over motion from the S-shaped two-dimensional wavelike motion. In turn, these movements increased the twisting force of the trunk, generating diversity and efficiency in the three-dimensional modes of movement. This locomotion pattern acquisition may partially recall the process by which modern terrestrial vertebrates evolved their walking behaviour.

The results of our experiments also suggest the possibility that sturgeons used a similar system to develop locomotion from reflex responses (Gesell et al., 1934; Karaca et al., 2013; McGraw, 1941). Fish such as mudskippers, Shuttles hoppfish, and blennies, which are known to thrive on land for extended periods, often move their tails quickly by bending them toward their heads and then straightening them, which allows the fish to move and jump (Ashley-Ross et al., 2013; Flammang et al., 2016; Hsieh, 2010; McFarlane et al., 2019; Nasuchon et al., 2016; Pace and Gibb, 2014) This common aspect in the locomotion of low-water-level-exposed sturgeon and fish that have achieved terrestrialisation, is attractive in terms of our understanding of the behavioural potential of fish.

A recent study reported that the tides on Earth 400 million years ago were stronger than present-day tides and that there is a correlation between strong tides and the places where the terrestrialisation of fish frequently occurred, and actinopterygian fossils were found (Balbus, 2014). In this study, we reared the sturgeons at low water levels for five days and underwater for two days, which allowed long-term rearing under low-water-level conditions. These conditions may be similar to the environment of 400 million years ago. Isotopic measurements and elemental analyses suggest that environmental changes such as hypoxia repeatedly occurred in the late Devonian period, implying that relatively large numbers of fishes may have used terrestrialisation as a passive escape behaviour depending on prevailing conditions (Joachimski and Buggisch, 2002; Murphy et al., 2000). Although primitive amphibians, such as Ichthyostega, are believed to have had large forelimbs and moved in a forelimb-driven manner on their belly, most animals including amphibians, and those that emerged after amphibians, developed their lower limbs for use in locomotion (BARCLAY, 1946; Pierce et al., 2012).

Regarding the functional and anatomical differences in the locomotion pattern, sturgeons took their posture with maximally spread pectoral fins when resting, whereas, during CSLC, the loads were applied to pectoral and pelvic fin regions on one side of their bodies with their fins folded. These results suggest that angle of pectoral girdle and fin was spread and the trunk-twisting action may lead to gradual increases in the contribution of the pelvic fin region in moving process on land. Although the sturgeons observed in this study do not represent dramatic species diversification, our results show the possibility that the longer a fish spends on land due to environmental changes, the greater the chances of gaining athletic benefits and ultimately may improve fish survival

## Acknowledgements

The authors wish to thank Dr. Atsushi Ishimatsu for his kind and helpful support in this study. The authors also wish to thank Dr. Haruo Hagiwara for his technical assistance. The Bester (hybrid sturgeons) used in this study were provided by the Fujikin Corporation. This work was supported by the Grant for Scientific Research Fund (No. 270002 and No. 300006) from the Teikyo University to A. Asano.

## Author contributions

Authors’ contributions Anshin Asano-Hoshino designed the study and wrote the initial draft of the manuscript. Drs. Hideyuki Tanaka, Nakakura Takashi, and Toshiaki Tsuji contributed to the analysis and interpretation of data, and Dr. Takuo, Mizukami, assisted in the preparation of the manuscript. All other authors have contributed to data collection and interpretation, and critically reviewed the manuscript. All authors approved the final version of the manuscript and agree to be accountable for all aspects of the work in ensuring that questions related to the accuracy or integrity of any part of the work are appropriately investigated and resolved.

## Supplementary information

**Figure S1.**
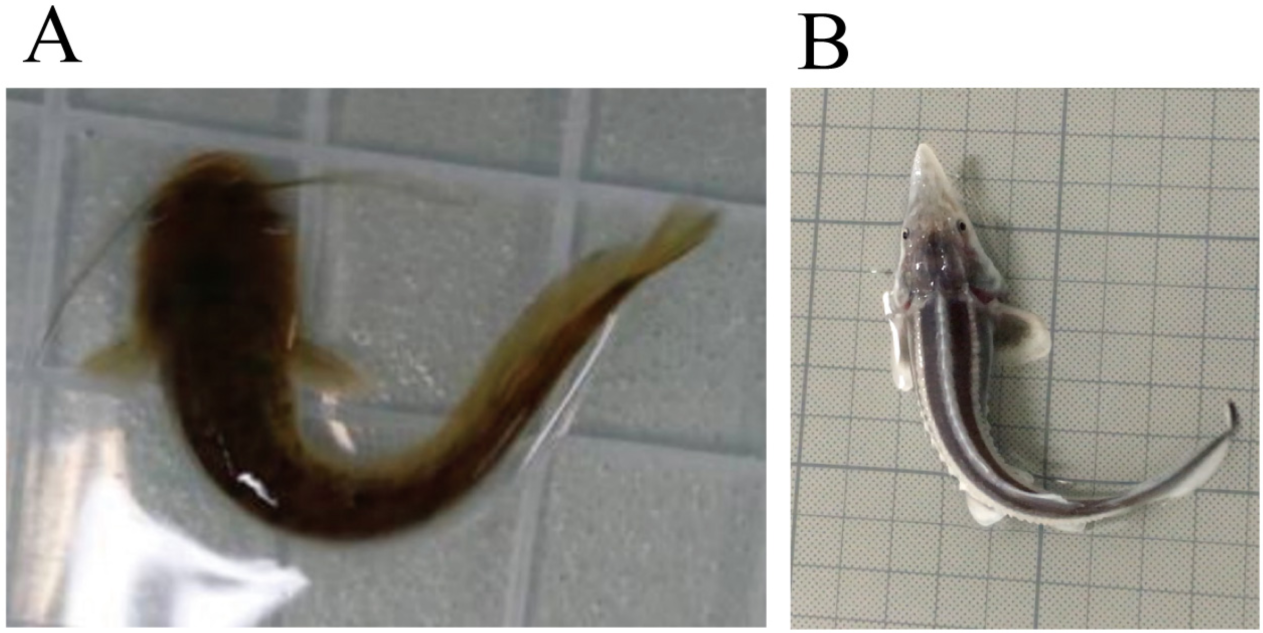
C-start escape response of catfish and juvenile sturgeons. (A) Catfish. (B) Juvenile sturgeon.

**Figure S2.**
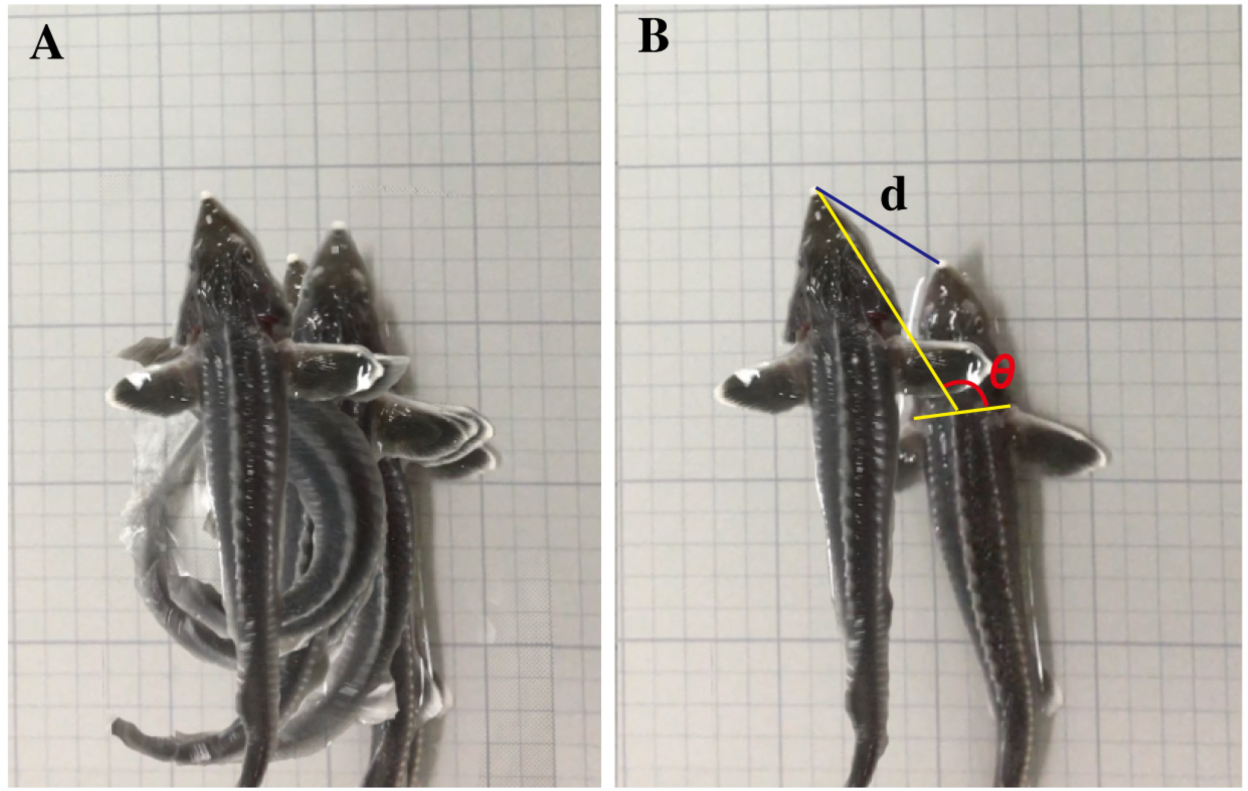
C-start escape response of sturgeons. (A) Overlapped sequential photographs showing the “C-start-like crawling” (CSLC) movement. (B).Start and end positions of CSLC. The black line (d) indicates the distance moved using CSLC. The yellow and red lines indicate the change in angle with CSLC (theta).

**Figure S3.**
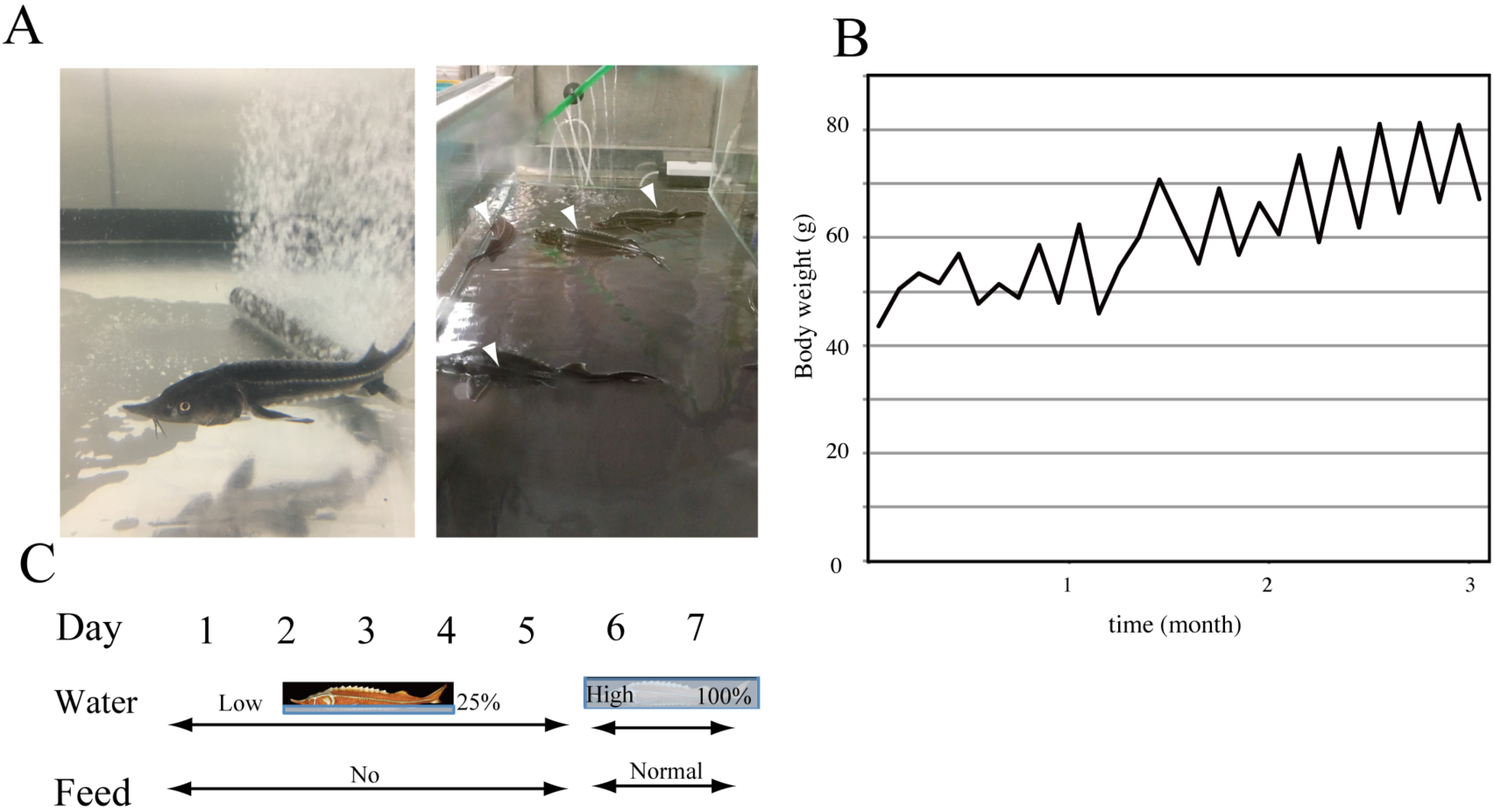
Changes in average body weight. (A) Sturgeon swimming underwater (left) and during exposure to low water levels (right). (B) Comparison between average body weight (g) and time (months); the average body weight of the sturgeon decreased during low-water-level exposure and increased during the return to higher water levels. (C) Schedule of the one-week rearing experiment.

**Figure S4.**
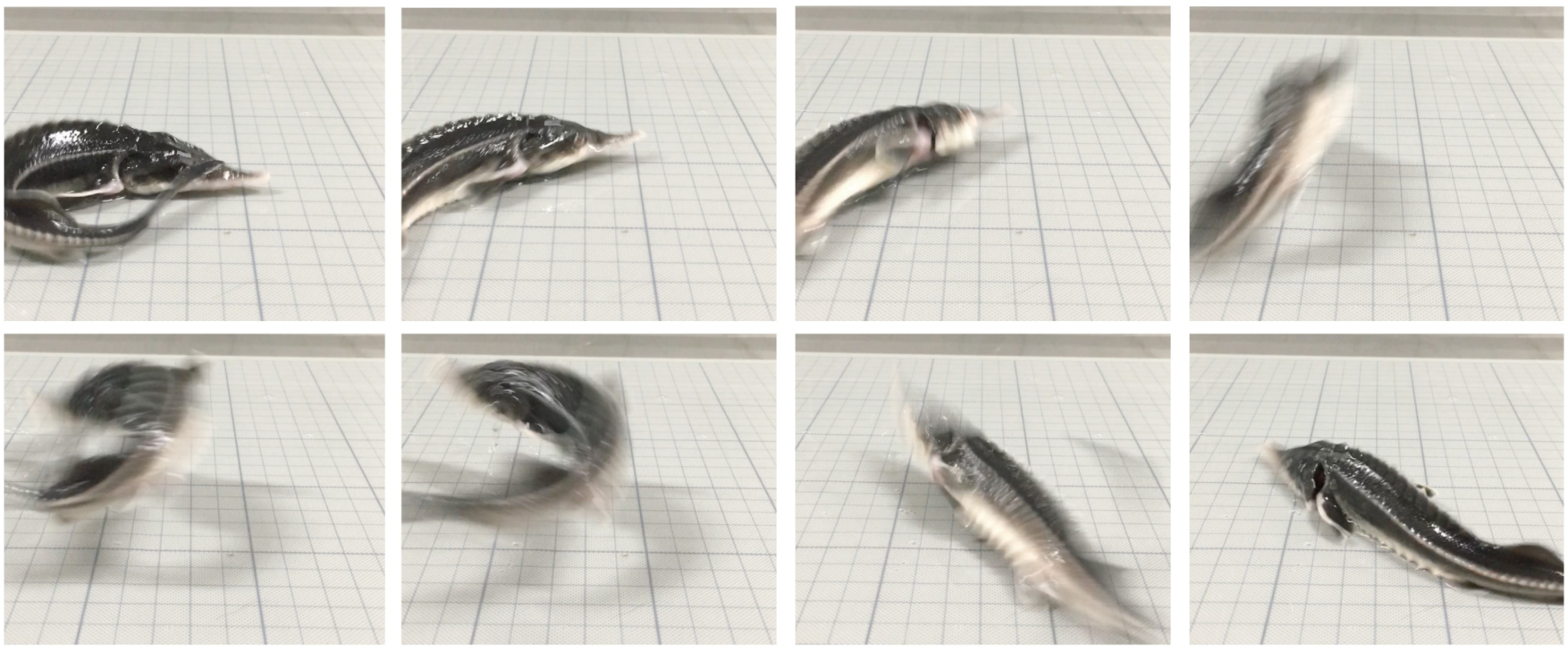
Sequential photographs of developmental locomotion.

**Movie S1**

The crawling movement and reflex response herein called the C-start escape response.

**Movie S2**

Sturgeon crawling with fully expanded pectoral fins.

